# Genomic epidemiology of a complex, multi-species plasmid-borne *bla*_KPC_ carbapenemase outbreak in Enterobacterales in the UK, 2009-2014

**DOI:** 10.1101/779538

**Authors:** N Stoesser, HTT Phan, Anna C. Seale, Zoie Aiken, Stephanie Thomas, Matthew Smith, David Wyllie, Ryan George, Robert Sebra, Amy J Mathers, Alison Vaughan, Tim EA Peto, Matthew J Ellington, Katie L Hopkins, Derrick W Crook, Alex Orlek, William Welfare, Julie Cawthorne, Cheryl Lenney, Andrew Dodgson, Neil Woodford, A Sarah Walker, the TRACE Investigators’ Group

**Author notes:** Corresponding author: Nicole Stoesser. These authors contributed equally to this work (N Stoesser, HTT Phan).

## Abstract

Carbapenem resistance in Enterobacterales is a public health threat. *Klebsiella pneumoniae* carbapenemase (encoded by alleles of the *bla*_KPC_ family) is one of the commonest transmissible carbapenem resistance mechanisms worldwide. The dissemination of *bla*_KPC_ has historically been associated with distinct *K. pneumoniae* lineages (clonal group 258 [CG258]), a particular plasmid family (pKpQIL), and a composite transposon (Tn*4401)*. In the UK, *bla*_KPC_ has caused a large-scale, persistent outbreak focused on hospitals in North-West England. This outbreak has evolved to be polyclonal and poly-species, but the genetic mechanisms underpinning this evolution have not been elucidated in detail; this study used short-read whole genome sequencing of 604 *bla*_KPC_-positive isolates (Illumina) and long-read assembly (PacBio)/polishing (Illumina) of 21 isolates for characterisation. We observed the dissemination of *bla*_KPC_ (predominantly *bla*_KPC-2_; 573/604 [95%] isolates) across eight species and more than 100 known sequence types. Although there was some variation at the transposon level (mostly Tn*4401*a, 584/604 (97%) isolates; predominantly with ATTGA-ATTGA target site duplications, 465/604 [77%] isolates), *bla*_KPC_ spread appears to have been supported by highly fluid, modular exchange of larger genetic segments amongst plasmid populations dominated by IncFIB (580/604 isolates), IncFII (545/604 isolates) and IncR replicons (252/604 isolates). The subset of reconstructed plasmid sequences also highlighted modular exchange amongst non-*bla*_KPC_ and *bla*_KPC_ plasmids, and the common presence of multiple replicons within *bla*_KPC_ plasmid structures (>60%). The substantial genomic plasticity observed has important implications for our understanding of the epidemiology of transmissible carbapenem resistance in Enterobacterales, for the implementation of adequate surveillance approaches, and for control.

**IMPORTANCE:** Antimicrobial resistance is a major threat to the management of infections, and resistance to carbapenems, one of the “last line” antibiotics available for managing drug-resistant infections, is a significant problem. This study used large-scale whole genome sequencing over a five-year period in the UK to highlight the complexity of genetic structures facilitating the spread of an important carbapenem resistance gene (*bla*_KPC_) amongst a number of bacterial species that cause disease in humans. In contrast to a recent pan-European study from 2012-2013(1), which demonstrated the major role of spread of clonal *bla*_KPC_-*Klebsiella pneumoniae* lineages in continental Europe, our study highlights the substantial plasticity in genetic mechanisms underpinning the dissemination of *bla*_KPC_. This genetic flux has important implications for: the surveillance of drug resistance (i.e. making surveillance more difficult); detection of outbreaks and tracking hospital transmission; generalizability of surveillance findings over time and for different regions; and for the implementation and evaluation of control interventions.

## INTRODUCTION

Antimicrobial resistance (AMR) in Enterobacterales is a critical public health threat. Carbapenem resistance is of particular concern, and outbreaks involving multiple species of carbapenemase-producing Enterobacterales (CPE) are increasingly reported(2-5). Exchange of AMR genes, including carbapenem resistance genes, happens at multiple genetic levels(6), and is often facilitated by their presence on plasmids [circular DNA structures of variable size (2kb∼>1Mb)], and/or other smaller mobile genetic elements (MGEs) such as transposons and insertion sequences (IS), that form part of the accessory genome.

Whole genome sequencing (WGS) has significantly improved our understanding of infectious diseases epidemiology and is used in both community-associated and nosocomial transmission analyses(7, 8). Although useful for delineating transmission routes in clonal, strain-based outbreaks, standard phylogenetic approaches and comparative analyses have been more difficult for outbreaks involving multiple bacterial strains/species and transmissible resistance genes(6). Reconstruction of the genetic structures of plasmids carrying relevant antimicrobial resistance genes using long-read sequencing has improved our understanding of the genetic complexity of these resistance gene outbreaks, but has been difficult to undertake on a large scale.

Although approximately 40 *Klebsiella pneumoniae* carbapenemase (KPC; encoded by *bla*_KPC_) variants have now been described (as per NCBI’s AMR reference gene catalogue, available at https://www.ncbi.nlm.nih.gov/pathogens/isolates#/refgene/), only two have been most widely reported globally, namely KPC-2 and KPC-3 (H272Y with respect to KPC-2; single nucleotide difference in *bla*_KPC_ [C814T])(9, 10). In the UK, the first KPC isolate identified was a KPC-4-containing *Enterobacter* sp. isolated in Scotland in 2003(11), with subsequent identification of KPC-3 in isolates in the UK in 2007. From 2007, increasing numbers of suspected KPC isolates were referred to Public Health England (PHE’s) Antimicrobial Resistance and Healthcare Associated Infections (AMRHAI) Reference Unit, with the majority of confirmed KPC-producers (>95%) coming from an evolving KPC-2-associated outbreak in hospitals in North-West England, first recognised in 2008(12). These isolates were predominantly *bla*_KPC_-positive Enterobacterales cultured from patients in the Central Manchester University Hospitals NHS Foundation Trust (CMFT; now part of Manchester University NHS Foundation Trust)(13). *bla*_KPC_ is thought to have been introduced into the region via a pKpQIL-like plasmid(14, 15), a plasmid backbone previously associated with the global dissemination of *bla*_KPC_ in *K. pneumoniae* clonal group 258, and already observed in other *K. pneumoniae* sequence types (STs) and species in an analysis of 44 UK KPC-Enterobacterales from 2008-2010(15).

We used WGS to undertake a large-scale retrospective study of this multi-species, polyclonal, *bla*_KPC_ outbreak in North-West England from 2009, generating complete genome structures, including *bla*_KPC_ plasmids, for a subset of isolates. We contextualised our analysis of regional outbreak strains using isolates from a national *bla*_KPC_ surveillance programme, with the goal of understanding the genetic structures associated with the regional emergence of *bla*_KPC_ in this setting.

## RESULTS

Of 742 isolates identified for sequencing, 60 (8%) were not retrievable or cultivable from the laboratory archives. After de-duplicating by taking the first *bla*_KPC_-positive Enterobacterales (KPC-E) per patient, and excluding sequencing failures, any sequences without *bla*_KPC_ (assumed lost in culture), and mixtures (identified from genomic data analysis, see Methods), 604 evaluable isolate sequences were included. These included: 327 archived isolates (54%) from inpatients in the early stages of the observed outbreak (2009-2011), of which 309 and 18 isolates were from CMFT and the University Hospital of South Manchester NHS Foundation Trust (UHSM; now part of Manchester University NHS Foundation Trust) respectively; 78 (13%) later isolates from CMFT/UHSM (2012-2014); 119 (20%) isolates from other hospitals (n=15 hospitals) in North-West England (2009-2014, excluding CMFT and UHSM, up to the first 25 consecutive KPC-E isolates per hospital); 72 (12%) isolates from UK and Republic of Ireland hospitals (n=72 locations [n=4 from the Republic of Ireland]) outside the North-West (2009-2014) (first KPC-E isolate per hospital); and 8 (1%) isolates from English outpatient/primary care settings.

Although three *bla*_KPC_ variants were observed in the 604 included isolates, *bla*_KPC-2_ dominated (n=573, 95%); *bla*_KPC-3_ [n=27, 4%] and *bla*_KPC-4_ [n=4, 1%]) were also observed. Two isolates (0.3%; trace524, trace534) showed evidence of mixed populations of *bla*_KPC-2_ and *bla*_KPC-3_. The median *bla*_KPC_ copy number estimate was 1.8 (IQR: 1.6-2.1), with a maximum of 8.2. Across the three main species, *bla*_KPC_ copy numbers were higher in *K. pneumoniae* (n=525 [87%], median 1.8 [IQR: 1.6-2.1]), than *E. coli* (40 [7%]: 1.7 [1.5-1.9]) or *E. cloacae* (26 [4%], 1.6 [1.4-2.0]) (Kruskal-Wallis; p=0.0003; Fig.1A). Amongst common STs, copy number was highest in *K. pneumoniae* ST258 (n=65 [11%], median 2.4 [IQR: 1.8-2.9]) versus other species/STs (n=531 [89%], median 1.8 [1.6, 2.0]) (Kruskal-Wallis; p=0.0001; Fig.1B, Fig.S1).

**Figure 1.**
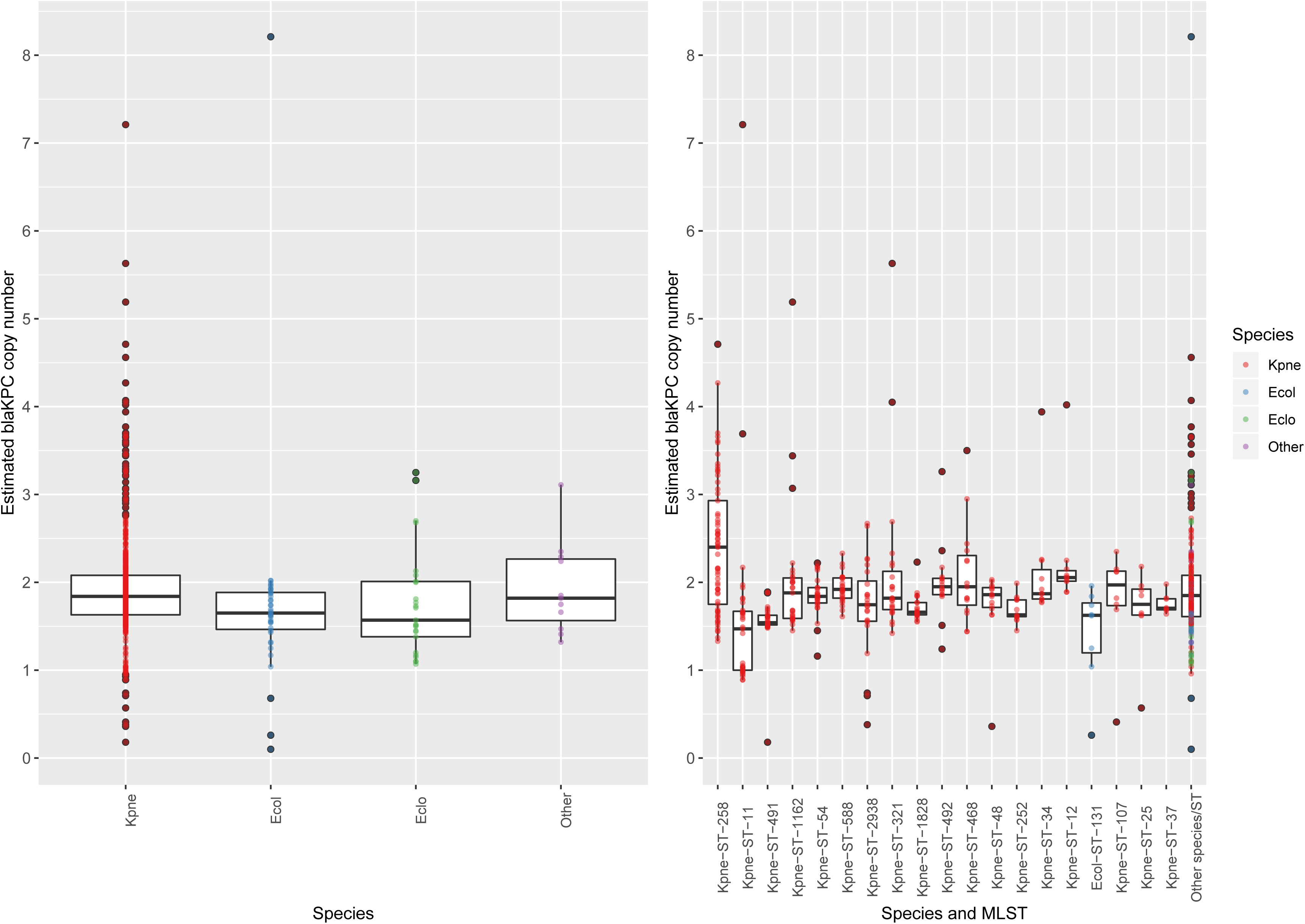
Estimated *bla*_KPC_ copy number distributions within major species (Fig.1A), and the top nineteen commonest species/ST combinations (Fig.1B) observed within the study (other ST/species combinations assigned as “Other” or “Other species/ST” respectively). Dots represent estimated copy number for single isolates; boxplots represent median estimated *bla*_KPC_ copy number +/- 1.58*IQR/sqrt(n). Boxplots are ordered by most common species and species/ST categories, left-to-right, except for the “Other”, “Other species/ST”, assigned to the right of the plots. For species assignations, “Kpne” = *Klebsiella pneumoniae*, “Ecol” = *Escherichia coli*, and “Eclo” = *Enterobacter cloacae*.

Other broad or extended-spectrum beta-lactamase genes were also commonly present across isolates, including: *bla*_TEM_ (n=452, all *bla*_TEM-1_), *bla*_OXA_ (n=492; Δ*bla*_OXA-9_ [n=425], *bla*_OXA-1_ [n=138]), *bla*_SHV_ (n=497) and *bla*_CTX-M_ (n=89; *bla*_CTX-M-15_ [n=57], *bla*_CTX-M-9_ [n=28]). Aminoglycoside resistance genes were also widely prevalent: *aac* (n=243), *aph* (n=196), *ant* (n=93) and *aadA* (n=280). In terms of acquired quinolone resistance, 160 isolates contained *qnr* variants, and 137 isolates contained *aac(6’)-Ib-cr*; no *qep* variants were seen.

Insertion sequences (ISs) have been shown to be key in the reshaping and streamlining of bacterial genomes, as well as exerting more subtle effects in the regulation of gene expression(16). The median number of different IS types in isolates was 15 (IQR: 13-16), with a maximum of 32. Four hundred distinct IS profiles were identified amongst the 604 isolates, with only five identical profiles shared amongst ≥10 isolates - these included a distinct profile seen only in national *K. pneumoniae* ST258 isolates (IS*1F*, IS*1R*, IS*3000*, IS*6100*, IS*903B*, IS*Ecl1*, IS*Kpn1*, IS*Kpn14*, IS*Kpn18*, IS*Kpn25*, IS*Kpn26*, IS*Kpn28*, IS*Kpn31*, IS*Kpn6*, IS*Kpn7*), and other unique profiles seen in small groups of *K. pneumoniae* ST588, ST11, ST321 and ST54. This highlights the significant flux of small mobile genetic elements within and between lineages in our dataset.

Tn*4401* is a ∼10kb transposon that has been the major transposable context for *bla*_KPC_ to date(17, 18). A predominant Tn*4401* isoform was associated with both *bla*_KPC-2_ and *bla*_KPC-3_ in this study, namely Tn*4401*a(17), which occurred in 584/604 (97%) isolates. Other known variants included Tn*4401*b (n=7) and Tn*4401*d (n=3). Only 20/584 (3%) isolates demonstrated evidence of SNV-level variation in Tn*4401*a (homozygous calls at 6 positions; heterozygous calls [i.e. mixed populations] at 3 positions). *bla*_KPC-2_-Tn*4401*a (n=539 isolates) was predominantly flanked by a 5-bp target site duplication (TSD) ATTGA, with 465/604 (77%; 465/539 [86%] of this sub-type) isolates with this Tn*4401*/TSD combination (Fig.2A). In 74 other *bla*_KPC-2_-Tn*4401*a isolates, the Tn*4401*a was flanked by other target site sequence (TSS) combinations, consistent with additional transposition events. Thirty-two of these were TSDs (16 AATAT-AATAT, 16 AGTTG-AGTTG), which have been described as more consistent with inter-plasmid transposition of Tn*4401*(19), and 35 were non-duplicate TSS combinations (ATTGA with either ATATA, TGGTA, CTGCC, AATAA, AGGAT), described as more consistent with intra-plasmid transposition. Evidence of multiple TSSs around *bla*_KPC-2_-Tn*4401*a within single isolates was seen in 6 cases (i.e. multiple right and/or left Tn*4401* TSSs); 1 case had a right TSS present, but no left TSS identified.

**Figure 2.**
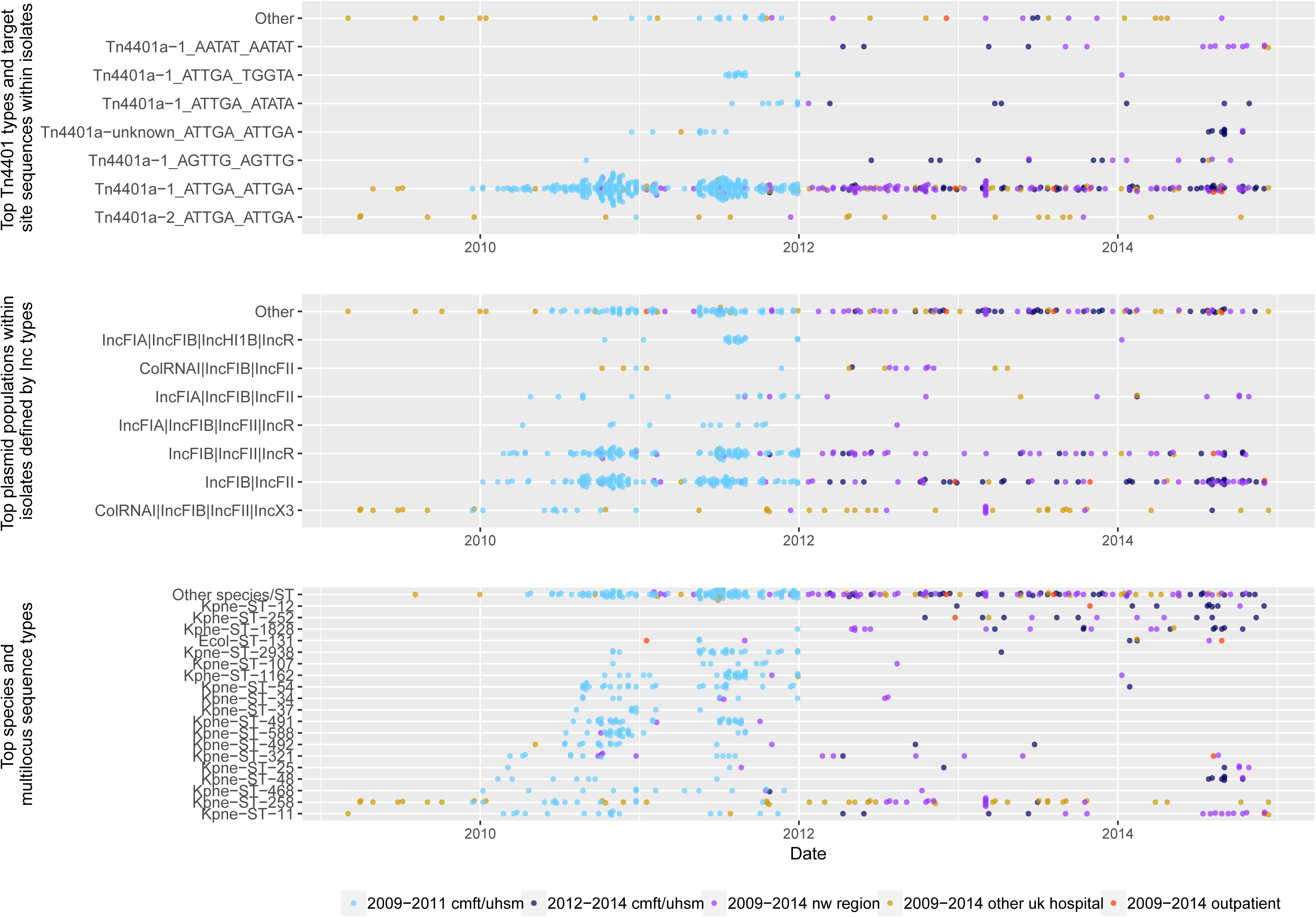
Incidence plots for 604 isolates included in the analysis. Dots are coloured by location of isolate collection, as defined in Methods. (A) Incidence plot of Tn*4401*/target site sequence (TSS; categories including ≥10 isolates); Tn*4401*a-1 is *bla*_KPC-2_/Tn*4401*a, Tn*4401*a-2 is *bla*_KPC-3_/Tn*4401*a; Tn*4401*a-unknown comprises a set of Tn*4401*a (n=18) with mutational variation including C684T, G962A C3042Y|G4739R, C4121T, G5583A and C7187T. (B) Incidence plot of plasmid replicon combinations (categories including ≥10 isolates). (C) Incidence plot of species/ST (top 20 common categories as in Fig.1).

The 604 isolates contained 91 unique combinations of plasmid Inc types (a crude proxy of plasmid populations present); no isolate was replicon negative. However, there were seven predominant combinations (Fig.2B) represented in 443/604 (73%) isolates, and these included six major Inc types, namely IncF (FIB [found in n=580 isolates], FII [n=545]), FIA (n=103), IncR (n=252), ColRNAI (n=86), and IncX3 (n=60). For many of the plasmid families, several different reference replicon sequences exist in the PlasmidFinder database, with a degree of homology amongst sequences in the same family, making it difficult to establish exactly which exact sub-type of replicon is present. However, restricting to 100% matches to reference replicon types for these common families, top matches included: IncFIB(pQil)_JN233705 (n=300) and IncFIB(K)_1_Kpn3_JN233704 (n=107); ColRNAI_1_DQ298019 (n=84); IncR_1_DQ449578 (n=70); and IncFII_1_pKP91_CP000966 (n=62; plasmid MLST IncFII_K4_), IncFII(K)_1_CP000648 (n=51; plasmid MLST IncFII_K1_) and IncFII_1_AY458016 (n=19; plasmid MLST IncFII_K2_).

Species and lineage diversity in the outbreak was substantial, with eight different species amongst sequenced isolates, and many different known STs, including: *K. pneumoniae* (n=525 isolates, 70 known STs), *E. coli* (n=40, 20 known STs), *Enterobacter cloacae* (n=26, 9 known STs), *Klebsiella oxytoca* (n=6, 3 known STs), *Raoultella ornithinolytica* (n=4), *Enterobacter aerogenes* (n=2), *Serratia marcescens* (n=1) and *Kluyvera ascorbata* (n=1). The most common STs were all *K. pneumoniae*, including ST258 (n=66), ST11 (n=35), ST491 (n=31), ST1162 (n=29) and ST54 (n=27) (Fig.2C). Therefore, although some of the earliest sequenced isolates were KPC-*K. pneumoniae* ST258 and ST11 (both in 2009) [two major KPC strains from CG258 circulating globally and in China at the time(9, 20)] and although KPC-producing *K. pneumoniae* ST258 appears to have been one of the earliest strains observed in CMFT and UHSM, multiple diverse STs and species were subsequently rapidly recruited to the outbreak in 2010 and 2011. This was most likely by the widespread sharing of a *bla*_KPC-2_-Tn*4401*a-ATTGA-ATTGA transposon within and between IncFIB, IncFII and IncR plasmid populations (Fig.2B, 2C).

### Long-read sequencing analyses

In addition to short-read data, to resolve genetic structures fully we obtained long-read PacBio data for 23 isolates, chosen to maximise the *bla*_KPC_ plasmid diversity assayed and focussing on isolates collected from the two main Manchester hospitals (12 CMFT isolates, 5 UHSM; plus 2 from other hospitals in North-West England, 4 from other UK locations). These included the two presumed earliest *bla*_KPC_ isolates from both CMFT and UHSM, as well as isolates sharing the same species/ST but with different plasmid replicon combinations or from North West regional versus national locations, same-species isolates with different STs, and isolates of different species. One PacBio sequencing dataset represented a clear isolate mixture (trace597 [UHSM] of *E. cloacae* ST133 and *K. pnemoniae* ST258), and for one isolate (trace457 [CMFT]), there were discrepancies between the short-read and long-read sequencing datasets, suggesting a laboratory error (*E. cloacae* ST45 long-read, *E. coli* ST88 short-read). These two assemblies were excluded, leaving 21 assemblies for further analysis (Table S1).

Of the 153 contigs from these 21 assemblies, 30 were clearly chromosomal, 77 plasmid, one chromosomal with an integrated plasmid, and 45 with unclear provenance (i.e. possibly phage, plasmid, or chromosomal). Overall 78/153 [51%] contigs were circularised, including 56/77 [73%] clear plasmid sequences. Thirty-one contigs (21 (68%) circularised) harboured *bla*_KPC_, of which one (trace552, *K. pneumoniae* ST11) had *bla*_KPC_ integrated into the chromosome. Four isolates had multiple copies of *bla*_KPC_ (trace205 [*K. pneumoniae*, ST468; 2 copies, 1 contig], trace457 [*E. cloacae*, ST45; 5 copies [1 truncated], 2 contigs], trace75 [*K. pneumoniae*, ST252; 2 copies, 2 contigs], trace149 [9 copies, 9 contigs]). Trace149 (*E. coli*, ST1642), which had nine copies of *bla*_KPC_, had one copy each on: a long, incomplete *bla*_KPC_ contig (203kb), seven nearly identical complete *bla*_KPC_-containing circularised sequences of size ∼10kb (possibly representing circularised translocatable intermediates(21)), and a short linear *bla*_KPC_ contig (∼18kb).

We observed *bla*_KPC_ in multiple plasmid backgrounds (Fig.3), including a majority of *bla*_KPC_ plasmids with multiple replicons (13/21 [60%] clear plasmid contigs, as represented in Fig.3), particularly with IncFIB/IncFII and/or IncR, consistent with replicon patterns in the isolates overall (Fig.2). For the IncFII group, for which we had 17 plasmid sequences with an IncFII(K)_CP000648-like replicon (plasmidFinder match; 5 *bla*_KPC_-negative [i.e. not represented in Fig.3] and 12 *bla*_KPC_-positive), there was evidence of significant exchange and rearrangement of plasmid components between both *bla*_KPC_-positive and *bla*_KPC_-negative plasmids, integration of IncFII_K_ and IncR plasmids, and gene duplication events of Tn*4401*/*bla*_KPC_, as well as sharing between STs and species (Fig.4).

**Figure 3.**
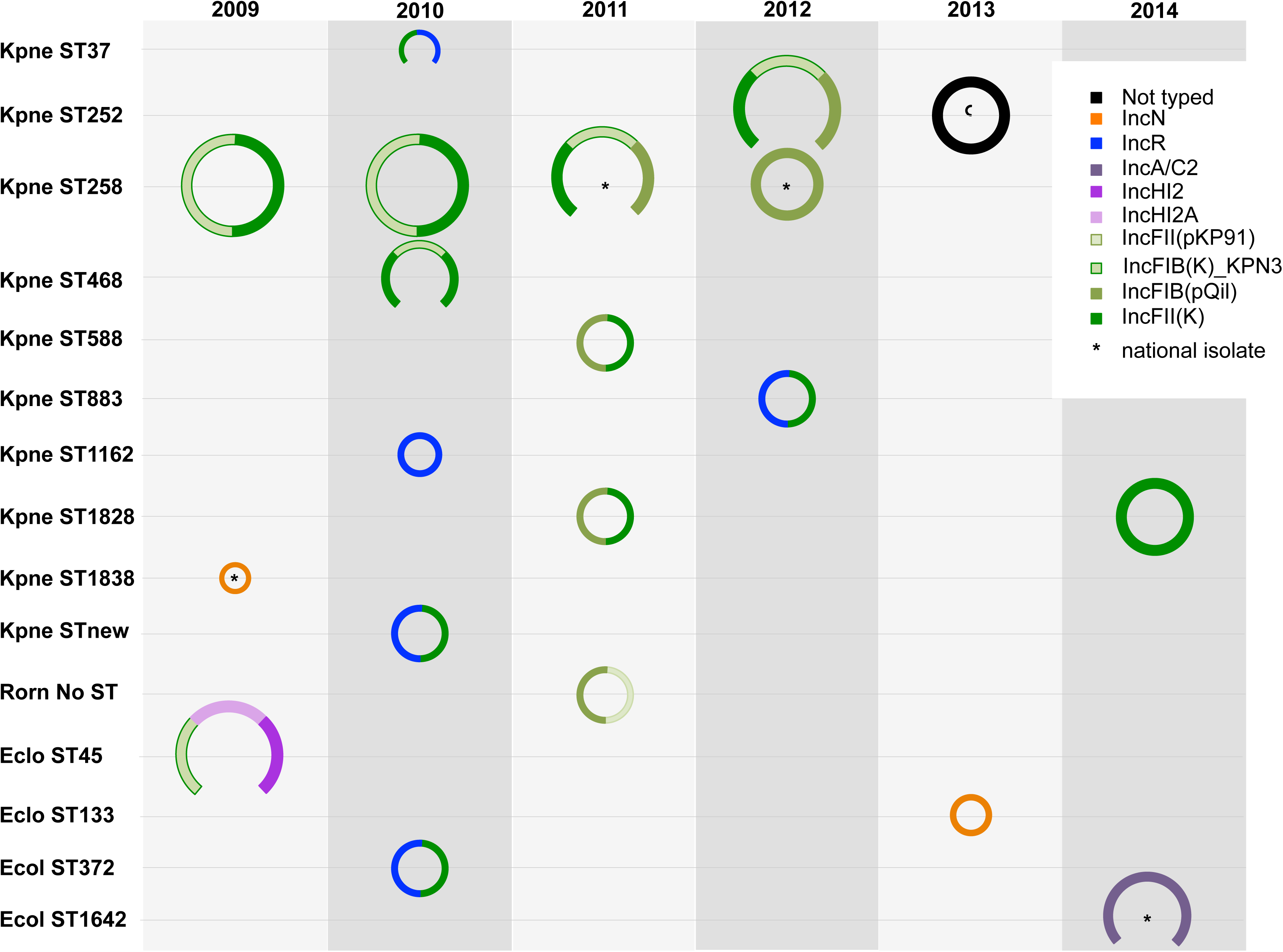
Schematic of *bla*_KPC_ plasmid types and sizes identified from long-read/short-read hybrid sequencing approach by species/ST and year of collection (NB only 21 contigs clearly designated as plasmid are represented). Closed circles denote circularised contigs (i.e. complete plasmids); circle colours denote replicon types assigned to each plasmid sequence (i.e. multiple colours represent multi-replicon plasmids). Plasmids from isolates from the wider UK collection are denoted with a “*”.

**Figure 4.**
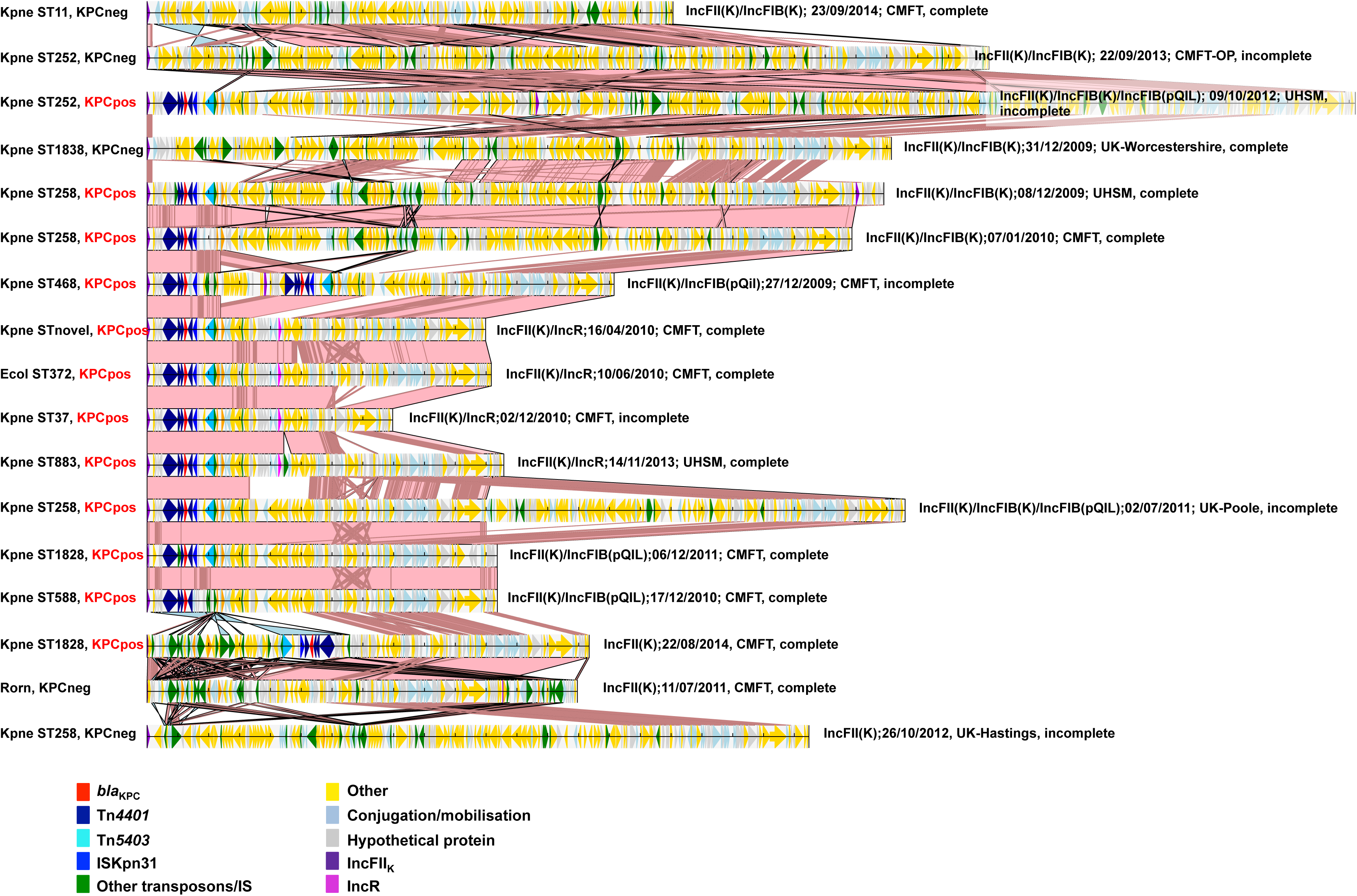
Alignments of plasmid sequences harbouring an IncFII(K)_1_CP000648-like replicon, includes *bla*_KPC_-negative and *bla*_KPC_-positive sequences. All sequences were re-orientated to start at IncFII for the purposes of alignment visualization (this also includes incomplete sequences, for which the exact structure and order may therefore be a proxy only). Loci of interest have been coloured and annotated as shown. Shading between sequences denotes regions of homology, with light pink shading denoting areas ≥90% nucleotide identity, dark pink areas ≥50% nucleotide identity, and light blue areas ≥90% nucleotide identity in reverse orientation. The order of sequences is adjusted to highlight genetic overlap between sequences, but not to imply any specific direct exchange events.

In addition to their plasticity, part of the success of these *bla*_KPC_ plasmids may also be attributable to the presence of toxin-antitoxin plasmid addiction systems (*ccdA*/*ccdB* n=4 *bla*_KPC_ plasmids; *higA* n=6; *vapB/vapC* n=11); anti-restriction mechanisms (*klcA* n=16, previously shown to promote *bla*_KPC_ dissemination(22)); and heavy metal resistance (*terB* [tellurite] n=3; *ars* operon [arsenicals] n=3; chromate resistance n=1; *cop* operon/*pcoC*/*pcoE* [copper] n=7; *mer* operon [mercury] n=10).

### bla_KPC_ plasmid typing

Attempts to identify complete plasmids (as opposed to plasmid replicon typing) from short-read data by comparison to a reference plasmid database has been estimated as being correct in only ∼45%-85% of cases in previous studies(6, 23). However, 13/14 (93%) of isolates for which we had hybrid assemblies with only one completely reconstructed *bla*_KPC_ plasmid had the correct top match using our *bla*_*KPC*_ plasmid typing method (Table S2). Noting that any complete plasmid typing approach from short-read data is sub-optimal, we compared all short-read sequences with our reference *bla*_KPC_ plasmid database (see Methods); matches to one or more reference *bla*_KPC_ plasmid sequences were identified in 554/604 (92%) isolates. Filtering the single match with the highest score at the predefined ≥0.80 threshold left a subset of 428/554 (77%) for evaluation. These 428 isolates had matches to 12 *bla*_KPC_ plasmid clusters.

Based on classification by these top plasmid-cluster matches, *bla*_KPC_ plasmid clusters were shared across a median (IQR) of 3 (1-6) STs, with pKpQIL-like plasmids being most widespread across species/STs (7 species, 75 STs), and clearly playing a major role in the North-West England outbreak, as well as being spread regionally and nationally (Fig.5). Other plasmid types identified as top-matches across the entire dataset included those fully resolved by long-read sequencing performed within this study, some of which were seen in ≥5% of study isolates (e.g. pKPC-trace75 [a non-typeable replicon]), and in non-North-West settings, likely reflecting recombination and generation of new *bla*_KPC_ plasmid variants in North-West England and their subsequent dissemination.

**Figure 5.**
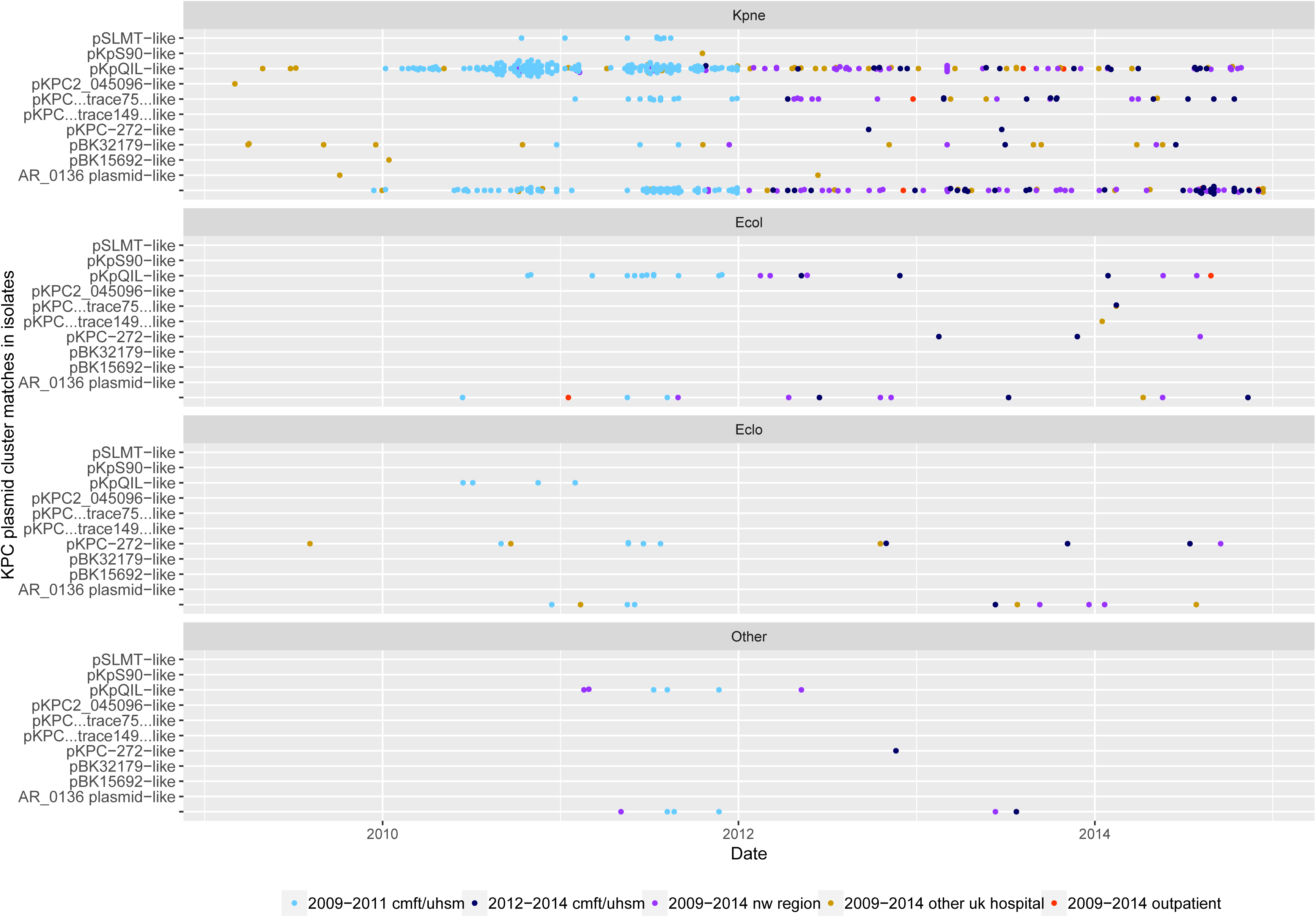
Short-read *bla*_KPC_ plasmid typing results (top match) for isolates by species and date. Dots are coloured by location of isolate collection, as defined in Methods.

## DISCUSSION

We present the largest WGS-based analysis of *bla*_KPC_-positive isolates (n=604) to our knowledge, focused on assessing genetic diversity around the carbapenemase gene itself rather than limiting the analysis based on species type, and incorporating a sampling frame from UK regional and national collections, over five years. *bla*_KPC_ remains one of the three most common carbapenemases observed in the UK, accounting for ∼15% of cases referred to the AMRHAI Reference Unit in 2017(24), and presenting a significant challenge to hospitals in North-West England, including Manchester, where it accounted for >97% of carbapenem resistance through 2015(25).

Our study provides an interesting context in which to consider the findings of a recently published pan-European survey of carbapenem non-susceptible *K. pneumoniae* (the EuSCAPE study; 6 months, 2013-2014; 244 hospitals, 32 countries)(1). In EuSCAPE, 684 carbapenemase-producing *Klebsiella* spp. isolates were Illumina sequenced, and similar to our study, most cases were healthcare-exposed (<2% from outpatients). EuSCAPE carbapenemase-producing isolates were also predominantly *bla*_KPC_ (∼45%, n=311 isolates), but mostly *bla*_KPC-3_ (232/311 [75%] versus 27/604 [5%] in our study), and ST258/ST512 (226/311 [73%] versus 107/525 (20%) of *K. pneumoniae* overall in our study). Based on identifying genetic “nearest-neighbours” in their data, the EuSCAPE team found 51% of *bla*_KPC_-*K. pneumoniae* were most closely related to another isolate from the same hospital. The authors concluded that there was strong geographic structuring of strains, and that the expansion of a handful of clonal lineages was predominantly responsible for the spread of carbapenemases in *K. pneumoniae* in Europe, with onward nosocomial transmission. Like *bla*_KPC-3_ in EuSCAPE, *bla*_KPC-2_ has also been linked with the clonal expansion of ST258 in Australia(26), where 48% of 176 *K. pneumoniae* isolates sequenced were *bla*_KPC-2_-containing ST258.

However, instead of clonal expansion, in our study we found rapid dissemination of mobile backgrounds supporting *bla*_KPC-2_, similar to observations from sequencing of other polyclonal *bla*_KPC_ outbreaks reported elsewhere, including the US(6, 27). Tn*4401*a, associated with high levels of *bla*_KPC_ expression(28), has been previously predominantly seen in *K. pneumoniae*, and in isolates from the US, Israel and Italy, and similarly most commonly with an ATTGA-ATTGA TSD(10). Thus our findings are consistent with the importation of the predominant *bla*_KPC-2_-Tn*4401*a-ATTGA-ATTGA motif into CMFT/North-West England and subsequent horizontal spread. Notably, as in EuSCAPE, 46/72 (64%) singleton isolates we sampled from UK hospitals were also CG258, but our detailed sampling within a region reflected a very different molecular epidemiology. Although the EuSCAPE study is large and impressive, its breadth may have been limiting in understanding regional diversity - for example, the subset of *bla*_KPC_-*K. pneumoniae* from the UK that were analysed in EuSCAPE consisted of 11 isolates submitted from six centres (https://microreact.org/project/EuSCAPE_UK). The focus was also more on analysing species-specific clonal relationships, with no analysis of other species or MGEs.

Although in our study diversification occurred at all genetic levels (Tn*4401*+TSSs, plasmids, plasmid populations, strains, species), there was more limited variation observed within the transposon and its flanking regions, and the spread of *bla*_KPC_ appears to have been supported by highly plastic modular exchange of larger genetic segments within a distinct plasmid population, particularly IncFIB/IncFII (found in 580 and 545 of the 604 isolates respectively) and IncR replicons (252/604 isolates). A previous study, in which 11 transformed *bla*_KPC_ plasmids from the UK (2008-2010) were sequenced (Roche 454/assembly, PCR+sequencing based gap closure), identified a UK variant of the pKpQIL plasmid, designated pKpQIL-UK (IncFII_K2_ by plasmid MLST), that was highly similar to pKpQIL (maximum 32 SNVs diversity), and several other IncFII_K2_ pKpQIL-like plasmids, but with novel segmental genetic rearrangements (gains/losses; pKpQIL-D1, pKpQIL-D2)(15). Our data support the importance of IncFII_K2_-like plasmids in this *bla*_KPC_ outbreak too, but also that other IncFII_K_-like plasmids (e.g. IncFII-_K1, -K4, -K7, -K15_) and replicons (IncFIB, IncR) have also been a significant feature. In addition to their plasticity, the plasmids identified frequently harboured AMR genes other than *bla*_KPC_ which might offer a selective advantage, alongside heavy metal resistance genes, and plasmid toxin-antitoxin addiction systems. The plasticity and association of IncFII_K_ plasmids with resistance genes and IncFIB replicons has been supported by findings of a recent analysis of IncFII_K_ plasmids(29).

The problem of accurately classifying plasmid populations from short-read data was exemplified in this analysis, and highlighted by our smaller long-read/short-read hybrid assembly-based analysis, which demonstrated significant diversity within structures assigned as similar by short-read based typing approaches. With this caveat, it was interesting that even with relatively relaxed thresholds, 29% of isolates did not have a match to our reference *bla*_KPC_ plasmid database (based on clustering of all publicly available reference sequences, as in Methods), consistent with rapid diversification in the genetic background of Tn*4401*/*bla*_KPC_ elements in this setting.

Our findings demonstrated that it is also important to consider plasmids without the resistance gene of interest in a population, as these may be relevant to a wider understanding of the transmission and evolution of smaller mobile genetic elements harbouring resistance genes (Fig.4). This was also shown to be relevant in a previous analysis of a large KPC-*E. coli* outbreak in the same setting in 2015-2016, in which a circulating *bla*_KPC_-negative plasmid, pCAD3 (IncFIB/FII), acquired Tn*4401* from a IncHI2/HI2A *bla*_KPC_-positive plasmid, and went on to dominate within a clonal *E. coli* lineage(25). Most studies in general however tend to focus on analysing AMR plasmids of interest. Fortunately, long-read sequencing is becoming increasingly low cost and high-throughput, and hybrid assembly is able to reconstruct plasmid sequences in Enterobacterales(30, 31). New developments in large-scale comparative genomics of complete genomes, including plasmid structures, are essential for future large-scale analysis of AMR gene outbreaks.

There are several limitations to our study. The reconstructed genomes generated using long-read PacBio data remained incomplete (49% of all contigs uncircularised). Improvements in long-read technology and assembly approaches will likely overcome this(30). Our short-read and long-read datasets were generated from the same frozen stocks of isolates, but from separate sub-cultures (because we used the short-read data to inform selection for long-read sequencing); ideally they would have been generated from the same DNA extract. PacBio sequencing library preparation incorporates size selection, and this may have led to short plasmid sequences (<15kb) being lost. Our interpretation of the evolution of backgrounds supporting *bla*_KPC_ was limited by the diversity present, and the inability to capture sequential evolutionary events, even with this large study. Lastly, very limited epidemiological data linked to the isolates were available, meaning that we were unable to ascertain any epidemiological drivers which might be contributing to the enormous heterogeneity of *bla*_KPC_ transmission over apparently short timeframes; the latter finding also precluded the useful application of standard phylogenetic approaches based on identifying variants core to and within species. In addition, the collection of isolates by PHE as part of regional and national surveillance was dictated by referral patterns of isolates from the hospitals surveyed, and we do not have any denominator information on cultures (either *bla*_KPC_-positive or *bla*_KPC_-negative) to corroborate details on the robustness of this referral process.

In conclusion, our large analysis highlights the difficulty and complexity of these outbreaks once important AMR genes have “escaped” the genetic confines of particular mobile genetic elements and bacterial species/lineages, with important implications for surveillance. These include the need to consider multiple bacterial species and plasmids as potential hosts of *bla*_KPC_, and invest resource in sequencing approaches to adequately reconstruct genetic structures and avoid misinterpreting the molecular epidemiology. It also demonstrates that regional differences in AMR gene epidemiology may be quite marked, which may affect the generalizability of control methods. Finally, it is important to consider the wider genetic background of host strains and plasmids in understanding the evolution and dissemination of important AMR genes, as AMR gene transfer between plasmid backgrounds within bacteria may occur over short timescales, and the interaction of several plasmids (i.e. not just those harbouring the AMR gene of interest at any given time) in a population may be highly relevant to the persistence and dissemination of the AMR gene itself.

## MATERIAL AND METHODS

### Study isolates and setting

We sequenced archived carbapenem-resistant Enterobacterales isolates from two hospital groups in Manchester (formerly known as CMFT and UHSM), aiming to include all inpatient isolates archived in the early stages of the observed outbreak, 2009-2011, and a subset of *bla*_KPC_-positive Enterobacterales (KPC-E) isolates sequenced as part of regional and national surveillance undertaken by Public Health England (PHE, 2009-2014). The PHE set included: (i) up to the first 25 consecutive KPC-E isolates from any hospital in North-West England (2009-2014) and referred to the PHE reference laboratory (2009-2014); (ii) the first KPC-E isolate from any other hospital in the UK and the Republic of Ireland referred to PHE (2009-2014); and, (iii) any KPC-E isolates from outpatient/primary care settings in the UK referred to PHE (2009-2014).

Ethical approval was not required as only bacterial isolates were sequenced, and their collection was part of outbreak investigation and management.

### DNA extraction and sequencing

For short-read Illumina sequencing (HiSeq 2500, 150bp PE reads), DNA was extracted using Quickgene (Fujifilm, Japan), with an additional mechanical lysis step following chemical lysis (FastPrep, MP Biomedicals, USA). Sequencing libraries were constructed using the NEBNext Ultra DNA Sample Prep Master Mix Kit (NEB) with minor modifications and a custom automated protocol on a Biomek FX (Beckman). Ligation of adapters was performed using Illumina Multiplex Adapters, and ligated libraries were size-selected using Ampure magnetic beads (Agencourt). Each library was PCR-enriched with custom primers (Index primer plus dual index PCR primer) (32). Enrichment and adapter extension of each preparation was obtained using 9ul of size-selected library in a 50ul PCR reaction. Reactions were then purified with Ampure beads (Agencourt/Beckman) on a Biomek NXp after 10 cycles of amplification (as per Illumina recommendations). Final size distributions of libraries were determined using a Tapestation 1DK system (Agilent/Lab901), and quantified by Qubit fluorometry (Thermofisher).

For long-read sequencing (PacBio [n=28]), DNA was extracted using the Qiagen Genomic tip 100/G kit (Qiagen, Netherlands). DNA extracts were initially sheared to an average length of 15kb using g-tubes, as specified by the manufacturer (Covaris). Sheared DNA was used in SMRTbell library preparation, as recommended by the manufacturer. Quantity and quality of the SMRTbell libraries were evaluated using the High Sensitivity dsDNA kit and Qubit Fluorimeter (Thermo Fisher Scientific) and DNA 12000 kit on the 2100 Bioanalyzer (Agilent). To obtain the longest possible SMRTbell libraries for sequencing (as recommended by the manufacturer), a further size selection step was performed using the PippinHT pulsed-field gel electrophoresis system (Sage Science), enriching for the SMRTbell libraries >15kb for loading onto the instrument. Sequencing primer and P6 polymerase were annealed and bound to the SMRTbell libraries, and each library was sequenced using a single SMRT cell on the PacBio RSII sequencing system.

Sequencing data have been deposited in the NCBI (BioProject Accession: PRJNA564424).

### Sequence data processing and assembly

We used Kraken(33) to assign species to sequenced isolates. SPAdes(34) v3.6 was used to *de novo* assemble reads (default options; subsequent removal of contigs shorter than 500bp and assembly coverage <2X). Isolates with sequence assemblies >6.5Mb were excluded to ensure that potentially mixed sequences were not included in the analyses. MLST was derived in silico by blasting de novo assemblies against publicly available MLST databases for *E. coli* (http://mlst.warwick.ac.uk/mlst/dbs/Ecoli), K. pneumoniae, E. cloacae and K. oxytoca (https://pubmlst.org/). Isolates with mixed MLST outputs were excluded.

Antimicrobial resistance (AMR) genes, plasmid replicon (Inc) types and insertion sequences (IS) were identified using resistType (https://github.com/hangphan/resistType_docker; curated AMR gene database as in(35), plasmid replicon reference sequences from PlasmidFinder(36), ISs from the ISFinder platform(37); ≥80% identity used as a threshold). *bla*_KPC_ copy number for each isolate was estimated by dividing coverage of the contig containing *bla*_KPC_ by the average coverage for the assembly. Plasmid MLST for common family types identified in short read data and for resolved genomes (i.e. based on PacBio and Pilon assemblies - see below) was confirmed by 100% sequence matches to reference alleles for families catalogued in the plasmidMLST website (https://pubmlst.org/plasmid/; IncA/C, IncHI1/2, IncN).

PacBio sequencing data were assembled using the HGAP pipeline(38), and polished with the corresponding Illumina datasets using Pilon (version 1.18, default parameters)(39). Chromosomal sequences and plasmid sequences were then manually curated where possible to create complete, closed, circular structures by using BLASTn to identify overlaps at the end of assembled contigs. Those with overlapping ends larger than 1000bp with sequence identity >99% were considered circularised/complete, and trimmed appropriately for resolution. Complete sequences were annotated using PROKKA (version 1.11)(40); annotations were used to determine genes known to encode toxin-antitoxin systems, heavy metal resistance, and anti-restriction mechanisms.

### Tn4401 typing

Tn*4401* typing was performed using TETyper(10), using the Tn*4401*, SNP and structural profile reference files included with the package (https://github.com/aesheppard/TETyper; Aug 2019), and a flanking length of 5bp, representative of the known target site signature sequence indicative of Tn*4401* transposition(41).

### *Plasmid database for bla*_KPC_ *plasmid typing*

A reference *bla*_KPC_ plasmid sequence database composed of *bla*_KPC_-harbouring contigs/plasmids from long-read sequencing of isolates in this study and all complete *bla*_KPC_ plasmids from (42-44) (August 2018) was used for *bla*_KPC_ plasmid typing within this study. To construct this database, all 279/6018 evaluable plasmid sequences carrying *bla*_KPC_ were first compared using *dnadiff*(45) to obtain the pairwise similarity between any two plasmid sequences *p*_*i*_ and *p*_*j*._ The similarity was defined as a function of their lengths *l*_*i*_, *l*_*j*,_ and the aligned bases *l*_*ij*_, *l*_*ji*_ as reported by:

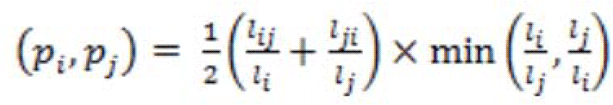

The score was designed to penalise differences in length of the compared sequences, i.e. to make sequences of different lengths proportionately more different. The resulting similarity matrix was used to perform clustering of plasmid sequences using the affinity propagation clustering technique, suitable for graph clustering problems with sparse similarity matrix and uneven cluster size and cluster number(46), and resulted in 34 clusters of 1-43 plasmids per cluster (Table S3). The largest cluster was the set of pKpQIL-like plasmids comprising 43 related sequences. Representative sequences of each *bla*_KPC_ plasmid cluster in this network were chosen randomly, to generate a set (*KPC-pDB*) of plasmids ranging from 7,995bp (NC_022345.1; plasmid pAP-2) to 447,095bp (NZ_CP029436.1; plasmid pKPC_CAV2013) in the final database used for *bla*_KPC_ plasmid typing in this study.

Subsequently, *bla*_KPC_ plasmid typing for each study isolate sequence was performed as follows: (1) assembled sequences for each isolate were BLASTed (BLASTn) against *KPC-pDB*; (2) any >1kb contig with >90% nucleotide identity and >80% total coverage match to sequences in *KPC-pDB* was retained; (3) for any sequence *p*_*i*_ in *KPC-pDB*, a score *s*_*i*_ was calculated by dividing the total matched bases of all contigs matched to *p*_*i*_ by *p*_*i*_’s length; and (4) an isolate was assumed to plausibly carry *p*_*i*_ if *s*_*i*_ ≥0.80. An isolate could have several *bla*_KPC_ plasmid matches; we restricted to the top match for each isolate in our analyses.

### Statistical analysis and data visualisation

Statistical analysis (Kruskal-Wallis testing) was carried out in Stata 14.2. Plots for figures 1, 2, 5 and S1 were generated using ggplot2 in R (version 1.1.463). Figure 4 was generated using the GenomeDiagram package(47) in Biopython(48).

## Acknowledgements and funding

We are grateful to and acknowledge the sharing of isolates by microbiology and clinical teams from contributing UK hospitals, and from Martin Cormican and the contributing laboratories in Dublin, Republic of Ireland. We are also grateful to the microbiology laboratory staff and infection control teams at Manchester University NHS Foundation Trust (formerly CMFT and UHSM); the staff of the Manchester Medical Microbiology Partnership; and the research laboratory, informatics and project management teams working as part of the Modernising Medical Microbiology consortium (Oxford).

Contemporaneous investigation by CMFT, UHSM and PHE was undertaken as part of routine activity. The retrospective investigation was funded by the National Institute for Health Research Health Protection Research Unit (NIHR HPRU) in Healthcare Associated Infections and Antimicrobial Resistance at Oxford University in partnership with Public Health England (PHE) [grant HPRU-2012-10041] and supported by the NIHR Biomedical Research Centre, Oxford. The views expressed in this publication are those of the authors and not necessarily those of the NHS, the National Institute for Health Research, the Department of Health or Public Health England. NS is funded by a PHE/University of Oxford Academic Clinical Lectureship. TEAP, DWC and ASW are NIHR Senior Investigators.

The Transmission of Carbapenemase-producing Enterobacteriaceae (TRACE) study investigators are listed alphabetically, and include several of the authors also listed by name in the main author list: Zoie Aiken, Oluwafemi Akinremi, Aiysha Ali, Julie Cawthorne, Paul Cleary, Derrick W. Crook, Valerie Decraene, Andrew Dodgson, Michel Doumith, Matthew J. Ellington, Ryan George, John Grimshaw, Malcolm Guiver, Robert Hill, Katie L. Hopkins, Rachel Jones, Cheryl Lenney, Amy J. Mathers, Ashley McEwan, Ginny Moore, Mark Neilson, Sarah Neilson, Tim E.A. Peto, Hang T.T. Phan, Mark Regan, Anna C. Seale, Nicole Stoesser, Jay Turner-Gardner, Vicky Watts, A. Sarah Walker, Jimmy Walker, David Wyllie, William Welfare and Neil Woodford.

None of the authors has any conflicts of interest to declare.

## Supplementary Tables and Figures

**Figure S1**. Estimated *bla*_KPC_ copy number distributions for species/ST combinations. Dots represent estimated copy number for single isolates; boxplots represent median estimated *bla*_KPC_ copy number +/- 1.58*IQR/sqrt(n). For species assignations, “Eclo” = *Enterobacter cloacae*, “Ecol” = *Escherichia coli*, “Ente” = *Enterobacter* spp., “Kluy” = *Kluvera* spp., “Koxy” = *Klebsiella oxytoca*, “Kpne” = *Klebsiella pneumoniae*, “Raou” = *Raoultella ornithinolytica*, “Serr” = *Serratia marcescens*.

**Table S1**. Details of isolates assembled using short-read (Illumina) and long-read (PacBio) datasets.

**Table S2**. Plasmid typing matches for isolates with short-read (Illumina) and long-read (PacBio) assemblies and reconstructed plasmid structures.

**Table S3**. Assignation of *bla*_KPC_ plasmids in study reference database to clusters for plasmid typing.

## References

1. David S, Reuter S, Harris SR, Glasner C, Feltwell T, Argimon S, Abudahab K, Goater R, Giani T, Errico G, Aspbury M, Sjunnebo S, Eu SWG, Group ES, Feil EJ, Rossolini GM, Aanensen DM, Grundmann H. 2019. Epidemic of carbapenem-resistant Klebsiella pneumoniae in Europe is driven by nosocomial spread. Nat Microbiol doi:10.1038/s41564-019-0492-8.

2. Mathers AJ, Stoesser N, Sheppard AE, Pankhurst L, Giess A, Yeh AJ, Didelot X, Turner SD, Sebra R, Kasarskis A, Peto T, Crook D, Sifri CD. 2015. Klebsiella pneumoniae Carbapenemase (KPC)-Producing K-pneumoniae at a Single Institution: Insights into Endemicity from Whole-Genome Sequencing. Antimicrobial Agents and Chemotherapy 59:1661-1668.

3. Cerqueira GC, Earl AM, Ernst CM, Grad YH, Dekker JP, Feldgarden M, Chapman SB, Reis-Cunha JL, Shea TP, Young S, Zeng Q, Delaney ML, Kim D, Peterson EM, O’Brien TF, Ferraro MJ, Hooper DC, Huang SS, Kirby JE, Onderdonk AB, Birren BW, Hung DT, Cosimi LA, Wortman JR, Murphy CI, Hanage WP. 2017. Multi-institute analysis of carbapenem resistance reveals remarkable diversity, unexplained mechanisms, and limited clonal outbreaks. Proc Natl Acad Sci U S A 114:1135-1140.

4. Ruiz-Garbajosa P, Curiao T, Tato M, Gijon D, Pintado V, Valverde A, Baquero F, Morosini MI, Coque TM, Canton R. 2013. Multiclonal dispersal of KPC genes following the emergence of non-ST258 KPC-producing Klebsiella pneumoniae clones in Madrid, Spain. J Antimicrob Chemother 68:2487-92.

5. Martin J, Phan H, Findlay J, Stoesser N, Pankhurst L, Navickaite I, De Maio N, Eyre D, Toogood G, Orsi N, Kirby A, Young N, Turton J, Hill R, Hopkins K, Woodford N, Peto T, Walker A, Crook D, Wilcox M. 2017. Covert dissemination of carbapenemase-producing Klebsiella pneumoniae (KPC) in a successfully controlled outbreak: long and short-read whole-genome sequencing demonstrate multiple genetic modes of transmission. Journal of Antimicrobial Chemotherapy.

6. Sheppard AE, Stoesser N, Wilson DJ, Sebra R, Kasarskis A, Anson LW, Giess A, Pankhurst LJ, Vaughan A, Grim CJ, Cox HL, Yeh AJ, Modernising Medical Microbiology Informatics G, Sifri CD, Walker AS, Peto TE, Crook DW, Mathers AJ. 2016. Nested Russian Doll-Like Genetic Mobility Drives Rapid Dissemination of the Carbapenem Resistance Gene blaKPC. Antimicrob Agents Chemother 60:3767-78.

7. Quainoo S, Coolen JPM, van Hijum S, Huynen MA, Melchers WJG, van Schaik W, Wertheim HFL. 2017. Whole-Genome Sequencing of Bacterial Pathogens: the Future of Nosocomial Outbreak Analysis. Clin Microbiol Rev 30:1015-1063.

8. Walker TM, Monk P, Smith EG, Peto TE. 2013. Contact investigations for outbreaks of Mycobacterium tuberculosis: advances through whole genome sequencing. Clin Microbiol Infect 19:796-802.

9. Munoz-Price LS, Poirel L, Bonomo RA, Schwaber MJ, Daikos GL, Cormican M, Cornaglia G, Garau J, Gniadkowski M, Hayden MK, Kumarasamy K, Livermore DM, Maya JJ, Nordmann P, Patel JB, Paterson DL, Pitout J, Villegas MV, Wang H, Woodford N, Quinn JP. 2013. Clinical epidemiology of the global expansion of Klebsiella pneumoniae carbapenemases. Lancet Infect Dis 13:785-96.

10. Sheppard AE, Stoesser N, German-Mesner I, Vegesana K, Sarah Walker A, Crook DW, Mathers AJ. 2018. TETyper: a bioinformatic pipeline for classifying variation and genetic contexts of transposable elements from short-read whole-genome sequencing data. Microb Genom 4.

11. Palepou MFW, N.; Hope, R.; Colman, M.; Glover, J.; Kaufmann, M.; Lafong, C.; Reynolds, R.; Livermore, D. M. 2005. Novel class A carbapenemase, KPC-4, in an Enterobacter isolate from Scotland, abstr abstr. 1134_01_20. Prog. Abstr. 15th Eur. Cong. Clin. Microbiol. Infect. Dis., Copenhagen, Denmark.

12. Public Health England. 2011. Carbapenemase-producing Enterobacteriaceae: laboratory confirmed cases, 2003 to 2013. https://www.gov.uk/government/publications/carbapenemase-producing-enterobacteriaceae-laboratory-confirmed-cases/carbapenemase-producing-enterobacteriaceae-laboratory-confirmed-cases-2003-to-2013. Accessed 02/09/2016.

13. Donker T, Henderson KL, Hopkins KL, Dodgson AR, Thomas S, Crook DW, Peto TEA, Johnson AP, Woodford N, Walker AS, Robotham JV. 2017. The relative importance of large problems far away versus small problems closer to home: insights into limiting the spread of antimicrobial resistance in England. BMC Med 15:86.

14. Findlay J, Hopkins KL, Doumith M, Meunier D, Wiuff C, Hill R, Pike R, Loy R, Mustafa N, Livermore DM, Woodford N. 2016. KPC enzymes in the UK: an analysis of the first 160 cases outside the North-West region. J Antimicrob Chemother 71:1199-206.

15. Doumith M, Findlay J, Hirani H, Hopkins KL, Livermore DM, Dodgson A, Woodford N. 2017. Major role of pKpQIL-like plasmids in the early dissemination of KPC-type carbapenemases in the UK. J Antimicrob Chemother 72:2241-2248.

16. Siguier P, Gourbeyre E, Chandler M. 2014. Bacterial insertion sequences: their genomic impact and diversity. FEMS Microbiol Rev 38:865-91.

17. Cuzon G, Naas T, Nordmann P. 2011. Functional characterization of Tn4401, a Tn3-based transposon involved in blaKPC gene mobilization. Antimicrob Agents Chemother 55:5370-3.

18. Cuzon G, Naas T, Truong H, Villegas MV, Wisell KT, Carmeli Y, Gales AC, Venezia SN, Quinn JP, Nordmann P. 2010. Worldwide diversity of Klebsiella pneumoniae that produce beta-lactamase blaKPC-2 gene. Emerg Infect Dis 16:1349-56.

19. He S, Hickman AB, Varani AM, Siguier P, Chandler M, Dekker JP, Dyda F. 2015. Insertion Sequence IS<em>26</em> Reorganizes Plasmids in Clinically Isolated Multidrug-Resistant Bacteria by Replicative Transposition. mBio 6:e00762-15.

20. Qi Y, Wei Z, Ji S, Du X, Shen P, Yu Y. 2011. ST11, the dominant clone of KPC-producing Klebsiella pneumoniae in China. J Antimicrob Chemother 66:307-12.

21. Harmer CJ, Hall RM. 2016. IS26-Mediated Formation of Transposons Carrying Antibiotic Resistance Genes. mSphere 1.

22. Liang W, Xie Y, Xiong W, Tang Y, Li G, Jiang X, Lu Y. 2017. Anti-Restriction Protein, KlcAHS, Promotes Dissemination of Carbapenem Resistance. Front Cell Infect Microbiol 7:150.

23. Arredondo-Alonso S, Willems RJ, van Schaik W, Schurch AC. 2017. On the (im)possibility of reconstructing plasmids from whole-genome short-read sequencing data. Microb Genom 3:e000128.

24. Public Health England. 2018. English Surveillance Programme for Antimicrobial Utilisation and Resistance (ESPAUR), Report 2018.

25. Decraene V, Phan HTT, George R, Wyllie DH, Akinremi O, Aiken Z, Cleary P, Dodgson A, Pankhurst L, Crook DW, Lenney C, Walker AS, Woodford N, Sebra R, Fath-Ordoubadi F, Mathers AJ, Seale AC, Guiver M, McEwan A, Watts V, Welfare W, Stoesser N, Cawthorne J, Group TI. 2018. A Large, Refractory Nosocomial Outbreak of Klebsiella pneumoniae Carbapenemase-Producing Escherichia coli Demonstrates Carbapenemase Gene Outbreaks Involving Sink Sites Require Novel Approaches to Infection Control. Antimicrob Agents Chemother 62.

26. Sherry NL, Lane CR, Kwong JC, Schultz M, Sait M, Stevens K, Ballard S, Goncalves da Silva A, Seemann T, Gorrie CL, Stinear TP, Williamson DA, Brett J, van Diemen A, Easton M, Howden BP. 2019. Genomics for Molecular Epidemiology and Detecting Transmission of Carbapenemase-Producing Enterobacterales in Victoria, Australia, 2012 to 2016. J Clin Microbiol 57.

27. Weingarten RA, Johnson RC, Conlan S, Ramsburg AM, Dekker JP, Lau AF, Khil P, Odom RT, Deming C, Park M, Thomas PJ, Program NCS, Henderson DK, Palmore TN, Segre JA, Frank KM. 2018. Genomic Analysis of Hospital Plumbing Reveals Diverse Reservoir of Bacterial Plasmids Conferring Carbapenem Resistance. MBio 9.

28. Cheruvanky A, Stoesser N, Sheppard AE, Crook DW, Hoffman PS, Weddle E, Carroll J, Sifri CD, Chai W, Barry K, Ramakrishnan G, Mathers AJ. 2017. Enhanced Klebsiella pneumoniae Carbapenemase Expression from a Novel Tn4401 Deletion. Antimicrob Agents Chemother 61.

29. Bi D, Zheng J, Li JJ, Sheng ZK, Zhu X, Ou HY, Li Q, Wei Q. 2018. In Silico Typing and Comparative Genomic Analysis of IncFIIK Plasmids and Insights into the Evolution of Replicons, Plasmid Backbones, and Resistance Determinant Profiles. Antimicrob Agents Chemother 62.

30. De Maio N, Shaw LP, Hubbard A, George S, Sanderson ND, Swann J, Wick R, AbuOun M, Stubberfield E, Hoosdally SJ, Crook DW, Peto TEA, Sheppard AE, Bailey MJ, Read DS, Anjum MF, Walker AS, Stoesser N, On Behalf Of The Rehab C. 2019. Comparison of long-read sequencing technologies in the hybrid assembly of complex bacterial genomes. Microb Genom doi:10.1099/mgen.0.000294.

31. Wick RR, Judd LM, Gorrie CL, Holt KE. 2017. Completing bacterial genome assemblies with multiplex MinION sequencing. Microb Genom 3:e000132.

32. Lamble S, Batty E, Attar M, Buck D, Bowden R, Lunter G, Crook D, El-Fahmawi B, Piazza P. 2013. Improved workflows for high throughput library preparation using the transposome-based nextera system. BMC Biotechnology 13:104.

33. Wood DE, Salzberg SL. 2014. Kraken: ultrafast metagenomic sequence classification using exact alignments. Genome Biol 15:R46.

34. Bankevich A, Nurk S, Antipov D, Gurevich AA, Dvorkin M, Kulikov AS, Lesin VM, Nikolenko SI, Pham S, Prjibelski AD, Pyshkin AV, Sirotkin AV, Vyahhi N, Tesler G, Alekseyev MA, Pevzner PA. 2012. SPAdes: a new genome assembly algorithm and its applications to single-cell sequencing. J Comput Biol 19:455-77.

35. Stoesser N, Batty EM, Eyre DW, Morgan M, Wyllie DH, Del Ojo Elias C, Johnson JR, Walker AS, Peto TE, Crook DW. 2013. Predicting antimicrobial susceptibilities for Escherichia coli and Klebsiella pneumoniae isolates using whole genomic sequence data. J Antimicrob Chemother 68:2234-44.

36. Carattoli A, Zankari E, Garcia-Fernandez A, Voldby Larsen M, Lund O, Villa L, Moller Aarestrup F, Hasman H. 2014. In silico detection and typing of plasmids using PlasmidFinder and plasmid multilocus sequence typing. Antimicrob Agents Chemother 58:3895-903.

37. Siguier P, Perochon J, Lestrade L, Mahillon J, Chandler M. 2006. ISfinder: the reference centre for bacterial insertion sequences. Nucleic Acids Res 34:D32-6.

38. Chin CS, Alexander DH, Marks P, Klammer AA, Drake J, Heiner C, Clum A, Copeland A, Huddleston J, Eichler EE, Turner SW, Korlach J. 2013. Nonhybrid, finished microbial genome assemblies from long-read SMRT sequencing data. Nat Methods 10:563-9.

39. Walker BJ, Abeel T, Shea T, Priest M, Abouelliel A, Sakthikumar S, Cuomo CA, Zeng Q, Wortman J, Young SK, Earl AM. 2014. Pilon: an integrated tool for comprehensive microbial variant detection and genome assembly improvement. PLoS One 9:e112963.

40. Seemann T. 2014. Prokka: rapid prokaryotic genome annotation. Bioinformatics 30:2068-9.

41. Naas T, Cuzon G, Villegas MV, Lartigue MF, Quinn JP, Nordmann P. 2008. Genetic structures at the origin of acquisition of the beta-lactamase bla KPC gene. Antimicrob Agents Chemother 52:1257-63.

42. Villa L, Feudi C, Fortini D, Brisse S, Passet V, Bonura C, Endimiani A, Mammina C, Ocampo AM, Jimenez JN, Doumith M, Woodford N, Hopkins K, Carattoli A. 2017. Diversity, virulence, and antimicrobial resistance of the KPC-producing Klebsiella pneumoniae ST307 clone. Microbial Genomics 3.

43. Orlek A, Phan H, Sheppard AE, Doumith M, Ellington M, Peto T, Crook D, Walker AS, Woodford N, Anjum MF, Stoesser N. 2017. A curated dataset of complete Enterobacteriaceae plasmids compiled from the NCBI nucleotide database. Data in Brief 12:423-426.

44. Stoesser N, Sheppard AE, Peirano G, Anson LW, Pankhurst L, Sebra R, Phan HTT, Kasarskis A, Mathers AJ, Peto TEA, Bradford P, Motyl MR, Walker AS, Crook DW, Pitout JD. 2017. Genomic epidemiology of global Klebsiella pneumoniae carbapenemase (KPC)-producing Escherichia coli. Sci Rep 7:5917.

45. Kurtz S, Phillippy A, Delcher AL, Smoot M, Shumway M, Antonescu C, Salzberg SL. 2004. Versatile and open software for comparing large genomes. Genome Biol 5:R12.

46. Frey BJ, Dueck D. 2007. Clustering by passing messages between data points. Science 315:972-6.

47. Pritchard L, White JA, Birch PR, Toth IK. 2006. GenomeDiagram: a python package for the visualization of large-scale genomic data. Bioinformatics 22:616-7.

48. Cock PJ, Antao T, Chang JT, Chapman BA, Cox CJ, Dalke A, Friedberg I, Hamelryck T, Kauff F, Wilczynski B, de Hoon MJ. 2009. Biopython: freely available Python tools for computational molecular biology and bioinformatics. Bioinformatics 25:1422-3.

