## Supplementary figures and images for "Genomic epidemiology of a complex, multi-species plasmid-borne *bla*_KPC_ carbapenemase outbreak in Enterobacterales in the UK, 2009-2014"

### Figure S1

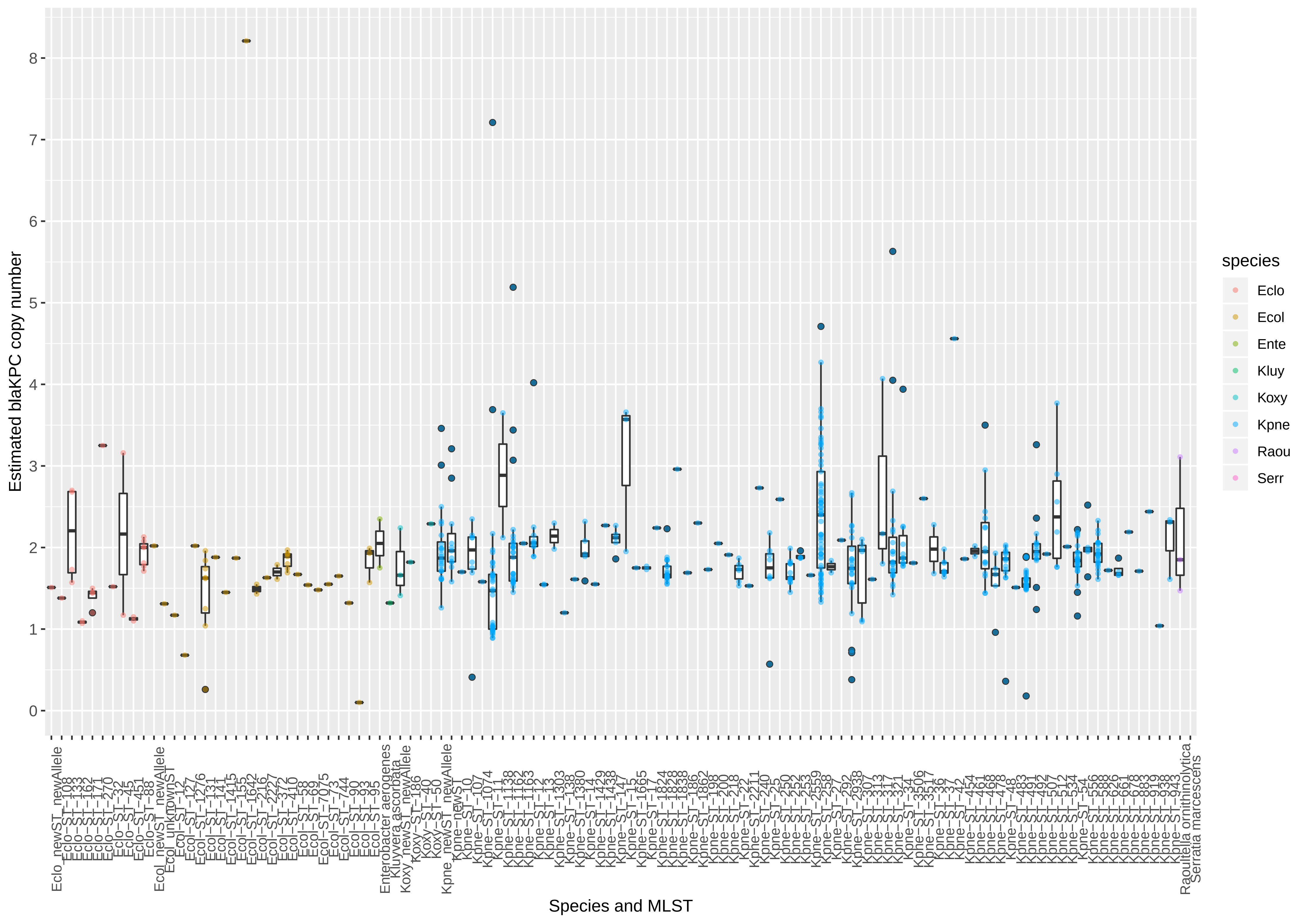
